# Experimental and Computational Evidence for Self-Assembly of Mitochondrial UCP2 in Lipid Bilayers

**DOI:** 10.1101/430835

**Authors:** A. Ardalan, S. O. Uwumarenogie, M. Fish, S. Sowlati-Hashjin, M. Karttunen, M. D. Smith, M. Jelokhani-Niaraki

## Abstract

Uncoupling proteins (UCPs) are members of the mitochondrial carrier family (MCF) that transport protons across the inner mitochondrial membrane, thereby uncoupling electron transport from ATP synthesis. The stoichiometry of UCPs, and the possibility of co-existence of this protein as monomeric and associated forms in lipid membranes remain an intriguing open question. In the current study, the tertiary structure of UCP2 was analyzed both experimentally and through molecular dynamics (MD) simulations. After recombinant expression of UCP2 in the inner membrane of *E. coli*, the protein was directly extracted from the bacterial membranes with a non-denaturing detergent and purified both as a pure monomer and as a mixture of monomers, dimers and tetramers. Both protein preparations were reconstituted in egg yolk lipid vesicles. Gel electrophoresis, circular dichroism spectroscopy and fluorescence methods were used to characterize the structure and the proton transport function of protein. UCP2 showed unique stable tetrameric forms in lipid bilayers. MD simulations using membrane lipids and principal component analysis support the experimental results and provided new molecular insights into the nature of noncovalent interactions in oligomeric UCP2. MD simulations indicate that UCP2 tetramers are asymmetric dimers of dimers, in which the interactions between the monomers forming the dimer are stronger than the interactions between the dimers within the tetramer. It is also shown that UCP2 has a specific tendency to form functional tetramers in lipid bilayers, capable of proton transport. The asymmetric nature of the UCP2 tetramer could act as a scaffold for regulating the activity of the monomeric units through cooperative intercommunication between these subunits. Under similar experimental conditions, the structurally comparable ADP/ATP carrier protein did not form tetramers in vesicles, implying that spontaneous tetramerization cannot be generalized to all MCF members.

**STATEMENT OF SIGNIFICANCE:** Self-assembly of membrane proteins plays a significant role in their biological function. In this article, both experimental and computational evidence are provided for spontaneous tetramerization of one of the mitochondrial uncoupling proteins (UCP2) in model lipid membranes. It is also shown that the tetrameric form of UCP2 is capable of proton transport, which leads to regulation of ATP synthesis in mitochondrion. Molecular dynamics simulations confirm the presence of asymmetric UCP2 tetramers as a potential scaffold for regulating the activity of the monomeric units through mutual intercommunication. The outcome of this study provides a solid ground for potential co-existence of monomeric and multimeric functional forms of UCPs that contributes to a deeper molecular insight into their structure and function.

## INTRODUCTION

Uncoupling proteins (UCPs) form a subfamily within the mitochondrial anion carrier superfamily (MCF), located in the inner membrane of mitochondria (IMM) (1). UCPs attenuate proton motive force by providing a back-flux route for protons to move from the intermembrane space (IMS) to the matrix, resulting in a decrease in the rate of ATP synthesis (2). Five human UCPs have been identified (UCPs 1-5). All of these five homologues transport protons and anions across the IMM and possess comparable, dominantly helical, conformations (3). UCP2 is unique among the UCPs in its ubiquitous expression in animal tissues (4). The wide distribution of UCP2 in various tissues suggests that it may play different roles under different physiological/pathological conditions (5). For example, a role for UCP2 in decreasing mitochondrial reactive oxygen species (ROS) concentration has been shown in several studies (6-8). Moreover, UCP2 appears to be capable of transporting C_4_ metabolites (9) and chloride ions (10). Up-regulation and anti-apoptotic properties of UCP2 in different cancerous cells have also been proposed (11-15).

Consistent with being a subfamily of the MCF superfamily, UCPs contain a tripartite structure composed of three tandem repeats. Each repeat comprises about 100 amino acids folded into two trans-membrane α-helices that are connected by a hydrophilic loop, positioned toward the matrix side of the mitochondrion (2, 16). Like other membrane proteins, isolation and purification of UCPs in their native functional states are challenging. All monomeric (17), dimeric (18, 19) and tetrameric (3) structures have been reported as the functional forms of UCPs. The monomeric structure of UCP1 is supported by electrophoretic and size exclusion assays and an isothermal titration calorimetry experiment in the decyl maltose neopentyl glycol detergent, which was also used (in combination with *n*-decyl-β-D-maltopyranoside) for the purification of the protein; however, despite their insistence on monomeric UCP1 as the only functional form of the protein, these researchers were not able to convincingly justify the appearance of higher molecular masses in their assays (17). The dimeric UCP structure has been a more traditional view evidenced by cross linking (19), ultracentrifuge (20) and nucleotide binding studies (21). The tetrameric form of UCP has been reported for recombinantly expressed proteins in bacterial membranes (not as inclusion bodies) and utilized circular dichroism spectroscopy (CD) analyses, as well as analytical ultracentrifuge and mass spectrometric methods for UCP1; this study did not exclude the possibility of co-existence of all three monomeric, dimeric and tetrameric UCPs as functional forms of the protein(s) (22). Based on the analysis of their contribution to mitochondrial Ca^2+^ uptake, it has also been suggested that UCPs co-assemble to form homo- (UCP2+UCP2) or hetero-multimers (UCP2+UCP3) (23). Moreover, assembly of UCPs with other MCFs, such as ADP/ATP carrier proteins, has been reported (24). Overall, the absence of a widely accepted high resolution structure for UCPs (25), variations in recombinant expression methodologies, and limitations accompanied by the choice and optimization of detergents for protein reconstitution in membranes are among the common shortcomings of all *in vitro* studies (22, 26).

Permanent and transient homo-oligomerization has been observed in many membrane proteins (27). Of note, among all enzymes listed in the Brenda enzyme database (https://www.brenda-enzymes.org), only 31.2% function as monomers and the rest (68.8%) are known to be oligomers (28). Self-assembly of proteins is not limited to soluble proteins; in fact co-assembly of integral membrane proteins to functional oligomers has been reported for several systems (29-32). It has been reported that proteins such as aquaporins and aquaglyceroporins (six membrane-spanning α-helices per subunit similar to that of UCPs) (29, 30), as well as several potassium channels from bacterial, archaeal or eukaryotic (31, 32) sources, can all form homo-tetrameric structures within the bilayer. Of homo- and hetero-tetrameric proteins many are, in fact, functional in the form of dimer of dimers (containing two different molecular interfaces; one between the monomers within the dimeric unit and another between the dimers). Examples of such proteins include streptavidin, transthyretin and hemoglobin (33). Association of membrane proteins (such as leucin transferase) is sometimes mediated and/or regulated by lipids, and delipidation by detergents can dissociate the multimeric proteins to monomers (27, 34). It has been shown that lipid molecules, such as cardiolipin, can also act as specific ligands to assist multimerization by bridging between monomers (34). The lipid membrane is therefore not only acts as a solvent for membrane proteins but also plays a crucial role in membrane proteins’ function, organization and assembly (35). Reconstitution of proteins into lipid membrane models, such as liposomes, is an essential tool in membrane protein research (36-38). Lipid constituents of liposomes can assist the self-association of subunits of oligomeric proteins within the bilayer (also addressed as the chaperoning effect) (31, 38). Self-assembly of bacterial potassium channel monomers to homo-tetramers in 1,2-dioleoylphosphatidylcholine (DOPC) liposomes is an example of lipid-assisted self-association (31); however, self-assembly of membrane proteins is not always lipid assisted. For reviews of lipid-protein interactions, see e.g., Refs. (39, 40).

In the current study, recombinantly produced UCP2 was reconstituted into lipid vesicles. Egg yolk PC has been chosen for making liposomes as it mainly contains PC (60%) and PE head groups, the most abundant lipids in mitochondrial membrane (41, 42). Protein’s conformation, function and self-association were investigated using CD, gel electrophoresis, western blotting and fluorescence spectroscopy. Moreover, the conformation and self-association behavior of UCP2 in lipid membranes was compared to the bovine ADP/ATP carrier protein isoform 1 (AAC1, PDB ID: 1OKC). This carrier protein is a member of the MCF with structural similarities to UCPs (1, 16, 43). Expression, purification and analyses techniques utilized for AAC1 were all similar/comparable to those of UCP2 to make the comparison logical. To further investigate the self-association of UCP2 in lipid membranes, atomistic molecular dynamics (MD) simulations were performed to complement the experimental methods. MD simulations have been shown to be a reliable approach to study the overall molecular conformations of membrane proteins and interactions determining membrane protein association in lipid membranes. See, for example Refs. (25, 44). Our experimental results, strongly supported by MD simulations, show that UCP2, specifically, forms stable and functional homo-tetramers (dimer of dimers) after reconstitution in phospholipid bilayers; in contrast, under the same conditions, AAC1 does not form tetramers in lipid bilayers and its mixed monomeric and dimeric form stays the same after reconstitution in lipid membranes.

## MATERIALS AND METHODS

### Chemicals

L-α-Phosphatidylcholine type X-E (L-α-PC) was from Sigma (St. Louis, Missouri). This mixed lipid system was extracted from dry egg yolk and contained more than 60% (by weight) phosphatidylcholine; the remaining lipids were mostly phosphatidylethanolamine and other phospholipids. C_8_E4 was from Bachem (Bubendorf, Switzerland). Octylglucopyranoside (OG) was obtained from BioVision (San Francisco, California). SPQ [6-methoxy-N-(3-sulfopropyl) quinoliniusm; 99% purity] was from Biotium (Fremont, California). Other chemicals were from Sigma-Aldrich.

### Overexpression and membrane extraction of proteins

Recombinant versions of UCP2 and AAC1 (UniProt Accession Codes UCP2 – P55851 and AAC1 – P02722, with poly-histidine affinity tags at their N terminal) were expressed in *E. coli* BL21 (DE3)-RIL (codon plus) and BL21 (DE3) cells using the auto-induction method as previously described (22, 45). Briefly, the bacterial cells were incubated for 22 hours in the auto-induction media [0.5% yeast extract, 0.5% glycerol, 1% tryptone, 0.2% lactose, 0.05% glucose, 50 mM KH_2_PO_4_, 50mM Na_2_HPO_4_, 1mM MgSO_4_, 25 mM (NH_4_)_2_SO_4_] at 22 °C. After 22 hours, the culture was centrifuged at 8000 g for 15 minutes and the bacterial cells (pellets) were collected. The pellets were resuspended in lysis buffer [500 mM NaCl, 20 mM Tris-HCl pH 8.0, one EDTA-free complete protease inhibitor tablet (Basel, Switzerland), 0.5 mg/ml DNase and 0.2 mg/ml lysozyme]. A high-pressure cell disruptor (Constant Systems Limited, Daventry, UK) was used to lyse the cells (at 20 kPsi). The lysate was centrifuged for 20 minutes at 20000 g to remove intact cells or aggregated proteins. The supernatant was collected and ultracentrifuged at 50000 g (MLA 80 rotor, Beckman Coulter) for 1 hour to obtain the bacterial membranes in the pellet. NADH oxidase activity assay was performed to verify the enriched presence of UCP in bacterial membranes (22). In this assay the membrane-embedded enzyme NADH catalytic activity is assessed based on its ability to convert NADH from its reduced to oxidized forms, as explained previously (22).

### Purification of proteins using immobilized metal affinity chromatography

Binding buffer containing 20 mM Tris-HCl, 1 mM THP [Tris (hydroxypropyl) phosphine], 500 mM NaCl, 1% LDAO (lauryldimethylamine oxide) detergent and 10 mM imidazole was used to solubilize the bacterial membrane. The solution was then incubated with Ni-NTA (Nickel-nitrilotriacetic acid, Thermo Scientific Waltham, Massachusetts) resin for 2 hours and subsequently transferred to a column. Detergent exchange (from 1% LDAO to 1% OG) was performed on the column at the washing step. In the case of UCP2 the resin in the column was washed with buffer containing 12 mM imidazole and successively eluted with buffers containing 25, 40 and 400 mM of imidazole in 1% OG detergent. The mixture of monomers and associated molecular forms of proteins were eluted at 40 mM imidazole and the pure monomer was eluted at 25 mM of imidazole. For AAC1, the column was washed with 30 mM imidazole and eluted with 400 mM imidazole in 1% OG detergent. Econo-Pac 10DG Columns (Bio-Rad, Hercules, California) were used to desalt the fractions in desalting buffer (20 mM Tris-HCl, 1% OG, 1% glycerol, pH 8.0). Purity and concentration of proteins were analyzed with semi-native PAGE (polyacrylamide gel electrophoresis) and modified Lowry assay (46), respectively. The desalted proteins were kept in desalting buffer and stored at −80° C.

### Semi-native PAGE analysis

To analyze proteins using semi-native PAGE (22), SDS was excluded from the sample buffer and was considerably reduced in the gel and running buffer as compared to SDS-PAGE conditions (2 mM for semi-native, compared to 35 mM in denaturing conditions). Moreover, samples were not heated before loading on the gel. The gels were run at 120 volts, stained with 0.2% w/v solution of Coomassie Brilliant Blue R-250 in acetic acid, methanol and water (10:45:45 by volume) for 2 hours and destained overnight.

### Western blot analysis

The identity of overexpressed recombinant wild-type UCP2 and AAC1 were confirmed by immunoblotting. Proteins (5-10 µg) were loaded on semi-native PAGE gels, and using the semi-dry technique, the proteins were transferred (75 minutes −110 volts) to nitrocellulose. The filters were stained with Amido Black to confirm transfer efficiency (47). The nitrocellulose filters were blocked for at least 60 minutes in Tris buffered saline containing 0.1% tween-20 and 5% dry skim milk. The primary antibody used to detect UCP2 was 1:2000 dilution of anti-UCP1/2/3 raised in rabbit (Santa Cruz Biotechnology Inc. Dallas, Texas), and for AAC1 the primary antibody was 1:5000 dilution of rabbit IgG anti-AAC1 (Bio-Rad). The secondary antibody was 1:5000 dilution of horseradish peroxidase (HRP) conjugated antibody raised in a goat against rabbit IgG (Rockland, Limerick, Pennsylvania). Luminata Crescendo Western HRP substrate (Millipore Sigma) was used as a chemiluminescent reagent to achieve immuno-detection. A Bio-Rad VersaDoc imaging system was employed to capture the image.

### Reconstitution in lipid vesicles

Reconstitution of UCP2 and AAC1 into lipid vesicles followed the procedure (3, 10) as described previously, with minor modifications. Briefly, solutions of egg yolk PC in MeOH: Chloroform (1:3) were dried for 8-12 hours under vacuum. The dried lipids were solubilized in an appropriate reconstitution buffer (supplemented with 3 mM SPQ (6-methoxy-1-(3-sulfonatopropyl) quinolinium) fluorescent probe in proton transport assays) to generate multilamellar vesicles which were consequently solubilized in C_8_E_4_ detergent to form lipid-detergent mixed micelles [at a lipid/detergent ratio of 1/2.5 (w/w)]. Pure proteins were added to the mixed micelles to reach the final 3 µM concentration. Protein/lipid ratio was ∼1/6000 (mol/mol) for proton transport assays and ∼1/600 (mol/mol) for CD experiments. Unilamellar proteoliposomes were spontaneously generated by removing the detergent using SM-2 Biobeads (Bio-Rad). The average radii of the prepared proteoliposomes for the proton transport assays and CD measurements were 76.87±2.29 nm and 31.46±1.68 nm, respectively, as determined by dynamic light scattering. On average, the radii of the blank liposomes were ∼13% larger than the proteoliposomes in both cases. In proton transport assays, part of the added SPQ was entrapped inside the liposomes and the external SPQ was removed by size-exclusion chromatography, using a Sephadex G25-300 (GE-Healthcare, Chicago, Illinois) spin column. Blank (protein-free) liposomes were prepared simultaneously in parallel with proteoliposomes.

### CD spectroscopy and fluorescence measurements

Far-UV CD measurements were performed at 25 °C, at 1 nm resolution, on an AVIV 215 spectropolarimeter (Aviv Biomedical, Lakewood, New Jersey). Quartz cells with 0.1 cm path length were used for measuring the CD spectra of the proteins in OG detergent and in liposomes. Ellipticities were converted to mean residue ellipticity, [θ]. Each spectrum was an average of 6-12 measurements.

Steady-state fluorescence of the liposomes was performed using a Cary Eclipse spectrophotometer (Varian, Palo Alto, California) with a band width slit of 5 nm and a scan speed of 600 nm/min. Excitation and emission of the SPQ fluorescence signal were at 347 and 442 nm, respectively. The fluorescence scans were performed at 25 °C. Each proton transport analysis was an average of 10 independent measurements.

### Proton transport measurements

Proton transport rates of the proteins were measured following their reconstitution in lipid vesicles, as described previously (10). Anions quench SPQ by a dynamic collision mechanism. In proton transport assays, 40 µL of liposomes were mixed with 1.96 mL of external buffer. The internal buffer was composed of TEA_2_SO_4_ (TEA: tetraethylammonium, 54 mM), TES buffer (20 mM) and EDTA (0.7 mM). The external buffer had a similar composition to the internal buffer, with TEA_2_SO_4_ replaced by K_2_SO_4_ (54 mM). The pH of both buffers was kept constant at 7.2. Palmitic acid (PA, final concentration 100 µM) was used to activate the UCP2-mediated proton flux. The addition of valinomycin mediated the influx of K^+^ which triggered fatty acid dependent proton efflux of UCP2. Proton efflux by UCP2 resulted in deprotonation of TES buffer. TES^−^ anions quenched the SPQ emission. The rate of decrease in the SPQ fluorescence signal was correlated to the rate of fatty acid activated proton transport of UCP2. The latter was measured by monitoring the decrease in SPQ’s fluorescence signal intensity within the first 30 seconds after addition of valinomycin (22). All proton transport data in liposomes were corrected for the basal non-specific leakage by subtraction of blank liposomes proton efflux. Furthermore, the proton transport data were calibrated for internal volume, SPQ’s fluorescence response and protein concentration. The concentration of protein in proteoliposomes was calculated using a modified Lowry assay (10, 46). No significant proton transport/leakage has been observed for the proteoliposomes in the absence of valinomycin. The inhibitory role of ATP on proton transport rates of reconstituted UCP2 was assessed by incubating the proteoliposomes with 500 µM ATP for 2 minutes prior for measurements.

### MD simulations

The only available high-resolution NMR structure of UCP2 (PDB ID: 2LCK) (16) was used as the starting structure for the monomeric UCP2. The same structure was used to build the dimeric and tetrameric forms. To form multimeric UCP2, the monomer units were aligned in such a way that the *GXXXG* (48), *GXXXAXXG* (49), and *SmXXXSm* (*Sm* = Gly, Ala, Ser, Thr) (50, 51) motifs were involved in the interactions with the neighboring unit(s) (Figure S1); these motifs have been proposed to promote helix-helix interaction (48). Moreover, in the case of dimer, several relative orientations for the monomers were examined. From these, the parallel orientation with maximal (distance-based) favorable interactions (electrostatic and hydrophobic) was chosen for the simulations. The tetramer was then built analogously from two dimers.

The protein-membrane systems were built using CHARMM-GUI (52). Membranes were built with 300, 316, and 498 1-palmitoyl-2-oleoylphosphatidylcholine (POPC) lipids for the monomeric, dimeric, and tetrameric UCP2, respectively. POPC was chosen to model the membranes since it is one of the main constituents of IMM (41, 53).

Each system was solvated in an explicit TIP3P water (54) box (total of 24195, 28463, and 47881 water molecules for monomer, dimer, and tetramer, respectively) and neutralized with 0.15 M KCl (65, 78, and 134 K^+^, and 80, 108, and 194 Cl^−^ for the monomer, dimer, and tetramer, respectively) to ensure overall charge neutrality. CHARMM36m/CHARMM36 protein/lipid force field were used (55, 56). All systems were first energy minimized using the steepest descents algorithm followed by 375 ps of preequilibration under constant volume and temperature (NVT), and constant pressure and temperature (NPT) conditions. MD simulations were conducted for 150 ns with 2 ps time steps using GROMACS/2018 (57). Chemical bonds were constrained using the linear constraint solver (P-LINCS) algorithm (58). The Nosé-Hoover thermostat (59, 60) was employed to maintain the temperature at 300 K with a 1.0 ps coupling constant and the Parrinello–Rahman barostat (61) with a compressibility of 4.5 × 10^−5^ bar^−1^ and a 5 ps coupling constant was used. The particle mesh Ewald method (62, 63) was used for the long-range interactions. The above protocol and methods have been shown to be reliable in a number of previous studies (e.g., Refs. (64-66) and references therein).

To study the dissociation process of the multimers, umbrella sampling (67) was conducted on the final structures obtained from the MD simulations. Selected windows of 0.1 nm along the pulling trajectories were first equilibrated for 100 ps and production runs were carried out for 5 ns. Protein subunits were pulled from their centers of masses until the subunits were separated by at least 1 layer of POPC. The weighted histogram analysis method (WHAM) was used for the analysis (68). Potential of mean force was monitored to confirm the end of the process. The bootstrap technique was used to estimate the statistical errors (69).

## RESULTS

### Expression and purification of UCP2 and AAC1

In order to study the quaternary structure of proteins, the cDNAs were cloned into the pET26b(+) expression vector and recombinant versions expressed in *E. coli* BL21 cells using the auto-induction method (45). Cloning the cDNAs of UCP2 and AAC1 into pET26b(+) resulted in the pelB signal peptide being added to the N-termini of the proteins, which targets the proteins to the *E. coli* inner membrane (3), followed by a a poly-histidine affinity tag. The pelB signal peptide normally directs secreted proteins to the periplasmic space; however, the transmembrane segments of UCP2 and AAC1 prevent full translocation and lead to insertion into the membrane due to hydrophobic interactions with the bacterial membrane (3, 22). Expression of proteins in the bacterial membranes allows the protein and its surrounding, tightly attached, membrane lipids to be extracted with mild detergents and co-purified. These co-purified membrane lipids can shield the protein from potential denaturing interactions with the solubilizing detergent, thus help UCP to remain relatively intact/folded. The His-tagged proteins were purified on nickel-containing columns and their purity was confirmed by semi-native PAGE (Figure 1). Bands of approximately 33, 66 and 132 kDa were observed for UCP2 on polyacrylamide semi-native gels. Furthermore, Western blot analysis using an α-UCP1,2,3 antibody reacted with all three bands (Figure 1) suggesting that they correlate with monomeric, dimeric and tetrameric forms of UCP2, respectively, as has been observed previously (3). In the case of AAC1, the monomeric and dimeric states were detected by semi-native PAGE (Figure 1) and Western blot analysis (Figure S2).

**Figure 1.**
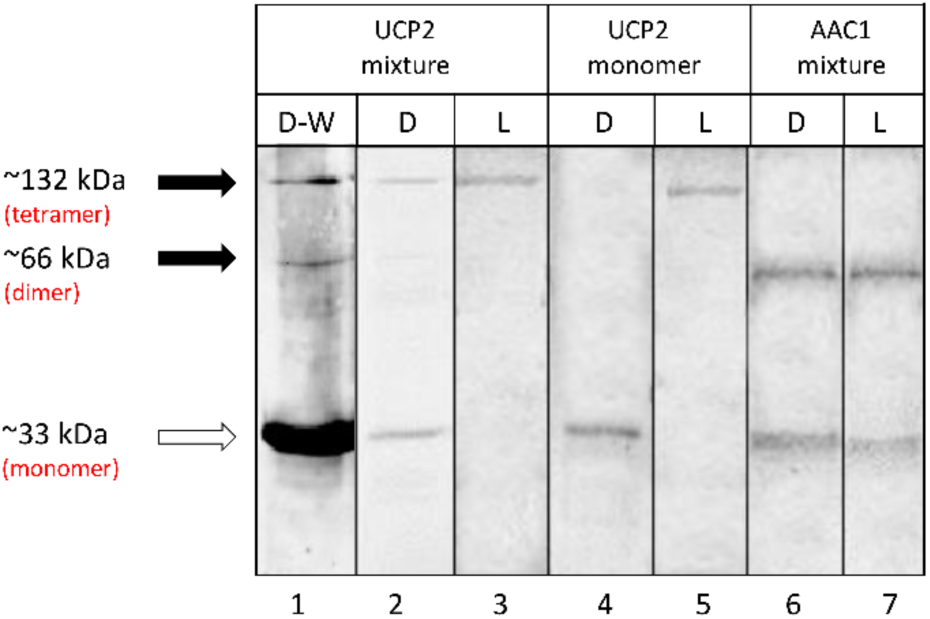
Semi-native PAGE analysis of purified UCP2 proteins in OG detergent (D) and in liposome (L) stained with Coomassie blue, and detected using Western blot (W) analysis. Mixture and monomer terms displayed at the top of the table refer to the protein that was used for reconstitution. The solid arrows show the location of oligomers and the open arrow shows the location of monomers. Lane 1 shows the Western blot detection of UCP2 in OG detergent probed with an α-UCP 1, 2, 3 antibody. Lanes 2-7 are stained with Coomassie. Collectively, the data confirm the presence of tetrameric UCP2 and the absence of tetrameric AAC1 in lipid vesicles. Lanes 3 and 5 correspond to proteoliposomes in which protein/lipid molar ratio was 1/6000.

### Semi-native PAGE analysis of UCP2 and AAC1 before and after reconstitution

As shown in Figure 1, UCP2 existed as a mix of monomeric, dimeric and tetrameric forms when purified in the presence of OG detergent, which has been chosen on the basis of our previous comparative CD and thermal stability analysis of the effect of different commonly used detergents [such as DDM (*n*-dodecyl-β-D-maltopyranoside), LDAO and OG] on UCP1 integrity. This analysis revealed that integrity/secondary structure of the protein was most stable when purified and stored in OG micelles (22). Once reconstituted in lipid vesicles from detergent, the resulting UCP2 proteoliposomes were analyzed using semi-native PAGE. As illustrated in Figure 1, only tetrameric forms of UCP2 are evident after reconstitution of the protein in liposomes (compare lanes 2 & 3).

To further investigate spontaneous oligomerization of UCP2 in lipid membranes, the protein was isolated from *E. coli* membranes in its monomeric form in the OG detergent and reconstituted in liposomes. The semi-native PAGE analysis of monomeric UCP2 before (in OG) and after reconstitution (in vesicles) is shown in Figure 1 (compare lanes 4 & 5). The semi-native gel results show that, under our experimental conditions, reconstitution of UCP2 monomers (in OG) into vesicles resulted in spontaneous self-association of UCP2 into a tetramer (Figures 1 and 2).

**Figure 2.**
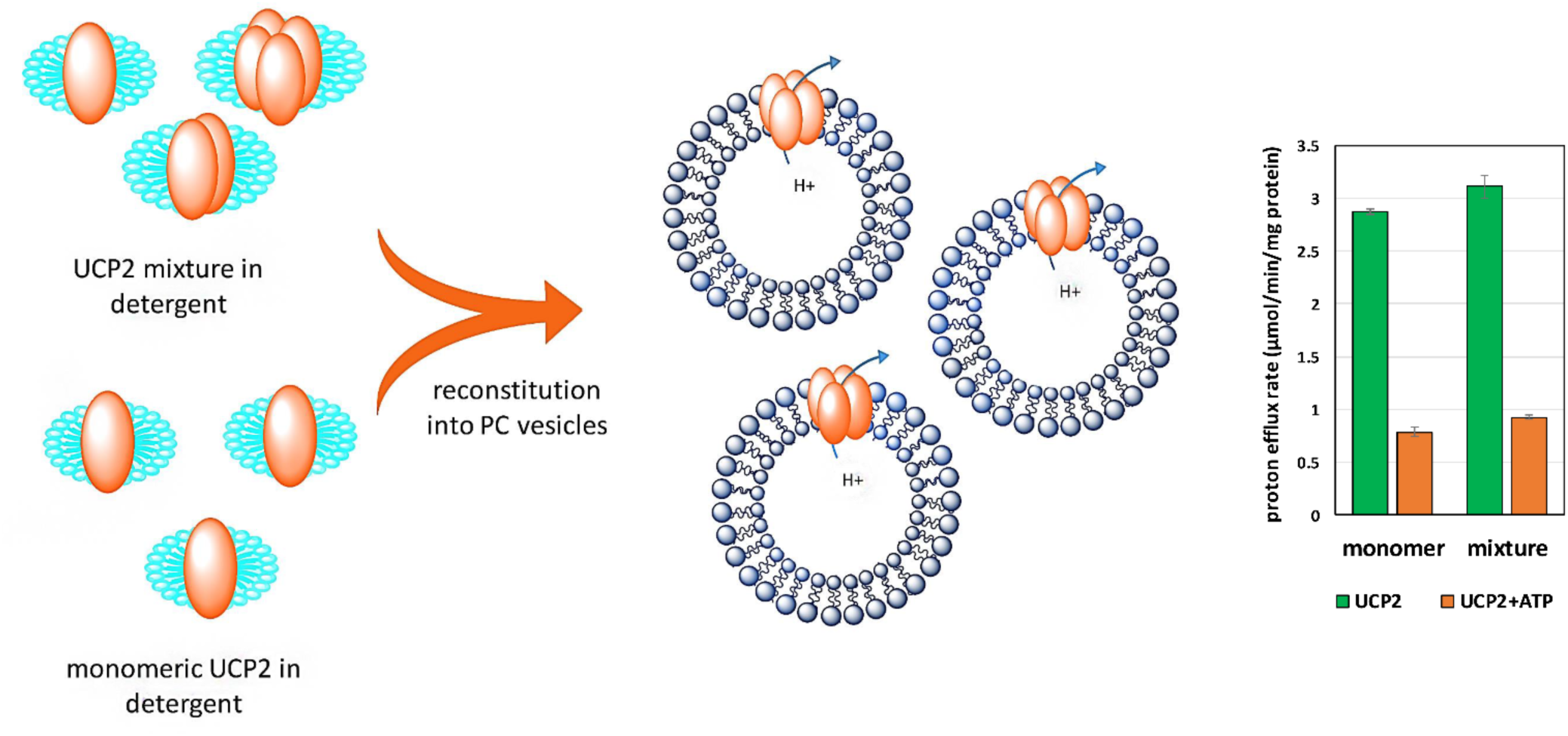
Formation of functional UCP2 tetramer after reconstitution in vesicles. Regardless of its molecular composition in detergent (monomeric or mixed monomeric/multimeric, before reconstitution), UCP2 adopts a tetrameric form in lipid vesicles. The bar graph on the right, shows the comparable rate of proton efflux for UCP2 in proteoliposomes generated either from monomeric or mixed molecular species. Each bar represents the average transport rate of 10 repeats and the error bars show the standard deviations.

In order to test the reliability of reconstitution and the probability of tetramerization of other MCF members, a mixture of purified AAC1 monomers and dimers were reconstituted into liposomes under the same experimental conditions as used for UCP2. The carrier protein AAC1 did not form tetrameric complexes in lipids; instead, it maintained its mixture of monomeric and dimeric forms (Figure 1, lanes 6 & 7). Other evidence for spontaneous self-association of UCP2 into homo-tetramers in liposomes comes from its proton transport function (Figure 2). Both reconstituted forms (from mixed and monomeric species in OG) of UCP2 resulted in similar ion transport profiles (Figure 2, right). This observation confirms the functional equivalence of reconstituted proteins from different preparations (monomeric vs. mixed monomeric/multimeric). The proton transport rates of tetrameric UCP2 resulted from reconstitution of monomeric and mixed monomeric/multimeric species in liposomes were comparable: 2.87 ± 0.028 and 3.11 ± 0.11 µmol/min/mg of protein, respectively. The proton transport function of the tetrameric proteins was further examined in the presence of a common inhibitor (ATP, 500 µM). Results showed that, in both cases, the rate of proton transport inhibition was ∼70% less compared to that of un-inhibited proteins (0.78±0.04 and 0.92±0.02 for proteoliposomes resulted from reconstitution of monomeric and mixed monomeric/multimeric proteins, respectively). These results exclude the possibility of non-specific proteoliposome leakage. The observed proton transport and inhibition rates are comparable with previous independent studies (3, 70, 71).

### CD spectroscopic conformational analyses of UCP2 and AAC1

CD spectroscopy was utilized to characterize and compare the conformations of UCP2 and AAC1 before (in OG) and after reconstitution into liposomes (Figure 3). The CD spectra of both proteins (UCP2 and AAC1) exhibited two negative maxima at 208 nm and 222 nm, as well as a positive maximum around 193 nm (not shown). Together, these spectral features are typical of a helical backbone structure of proteins, corresponding to π → π* (193 and 208 nm) and n → π* (222 nm) transitions in the peptide bond (72).

**Figure 3.**
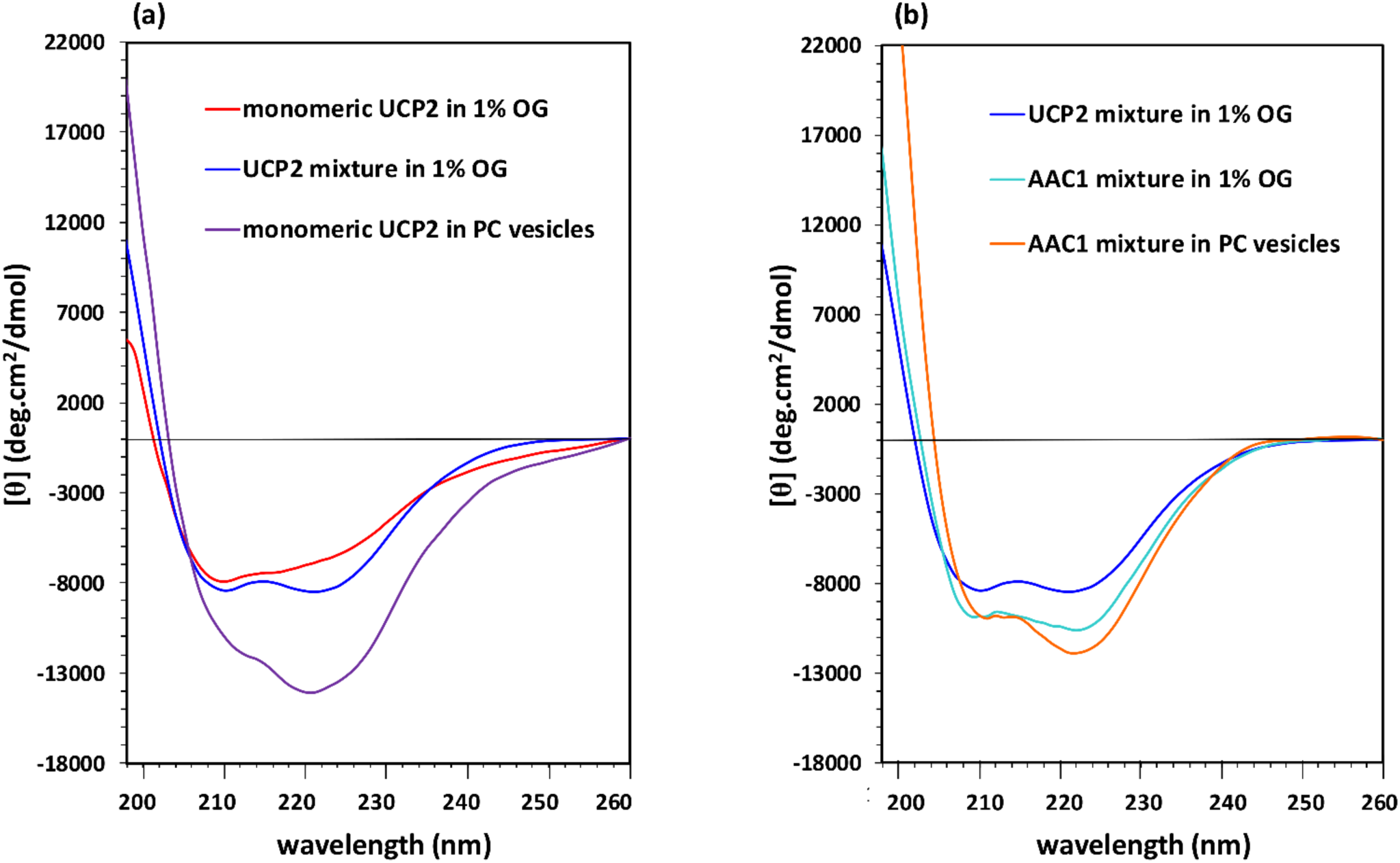
CD spectra of purified UCP2 and AAC1 before (in OG detergent) and after reconstitution in liposomes at 25 °C. (a) Tetramer formation of UCP2, regardless of its original molecular form in detergent, appeared as a conspicuous change in CD spectrum. (b) Comparable CD spectra of AAC1 before and after reconstitution implies a comparable molecular composition of the protein in liposome *vs* detergent, which is also similar to the spectrum of the UCP2 mixture in detergent. Concentrations of proteins were 5 – 8 µM in OG and ∼ 1 µM in lipid vesicles.

In UCP2, the far-UV CD signal was notably enhanced after reconstitution into liposomes, indicating a higher helicity compared to that of protein in OG detergent (Figure 3a). This marked conformational change, reflected in enhancement of negative ellipticity and reversal of the relative intensity of minima, reveals the important role of the lipid environment in the folding and structural stabilization of UCP2. In addition to their difference in signal intensity, the shape of the CD spectrum of UCP2 in proteoliposomes differed from that of the monomer (and the mixed monomer/multimer) in OG. Particularly, in lipid vesicles, a shoulder-like π **→** π* parallel transition band around 208 nm replaced the minimum band in OG, and also a considerably more intense n **→** π* transition band appeared at 222 nm. The θ_208_/θ_222_ ellipticity ratio of UCP2 monomer spectrum changed from 1.07 to 0.68 after reconstitution in liposomes (Figure 3a). The θ_208_/θ_222_ ellipticity ratios lower than one has been previously reported for the UCP helical bundle motifs and self-associated proteins (3). Relative decrease in molar ellipticity at 208 nm (in comparison to ellipticity at 222 nm) and its red shift, due to intramolecular interactions between transmembrane (TM) helical motifs and intermolecular protein association, have been also reported for UCP1 and other membrane proteins (22, 73-75). The overall pattern of the CD spectra (Figure 3a) related to the tetrameric form of UCP2 is in good agreement with our previously reported studies on the molecular forms of reconstituted UCPs in liposomes (3, 22). The CD spectra of the mixed molecular forms of UCP2 (monomer, dimer and tetramer) and AAC1 (monomer and dimer) in OG detergent exhibited comparable helical conformations (Figure 3b); as expected, based on the known similarities of the structures and co-presence of different monomeric and associated molecular forms in both proteins.

In comparison with UCP2, the negative ellipticity for AAC1 in liposomes was less enhanced (Figure 3b). The θ_208_/θ_222_ ellipticity ratio of AAC1 changed from 0.89 in OG detergent to 0.84 after reconstitution in liposomes, which is consistent with the protein maintaining similar conformations in these environments (mix of monomeric and dimeric forms) (Figure 3b). The differences between the CD spectra of reconstituted UCP2 and AAC1 show that the two proteins acquire different molecular forms after reconstitution in vesicles. In liposomes, both the lower negative ellipticity and higher θ_208_/θ_222_ ellipticity ratio of AAC1 compared to UCP2 (0.84 vs. 0.68) are consistent with a lower degree of association in AAC1.

In summary, our results from gel electrophoresis, CD spectroscopy and proton transport assays suggest that the structurally comparable UCP2 and AAC1 behave differently in liposomes. UCP2 associates into a functional tetramer regardless of its oligomeric state prior to reconstitution, while AAC1 retains its oligomeric state after reconstitution. Furthermore, only monomeric and/or dimeric forms (but no tetrameric forms, as found in UCP2) were detectable for AAC1.

### MD simulations

It has been well established that interactions of TM helices within the lipid bilayers have crucial roles in folding (76), stability (77) and function (78) of membrane proteins. To obtain atomic-level details of the protein-protein and protein-lipid interactions and to further evaluate our experimental results, all-atom MD simulations were conducted on monomeric, dimeric and tetrameric UCP2 in POPC model bilayers. This phospholipid is one of the main constituents of the IMM (41, 53). Moreover, umbrella sampling simulations were carried out to examine the stability of the multimeric forms and to provide further insights into the protein’s subunit interactions and multimerization process.

The general structural features of the protein were preserved in all three UCP2 systems through intensive interactions with the phospholipid bilayer (Figure 4). The average root-mean-square-deviations (RMSDs) of the protein backbone and the root-mean-square-fluctuations (RMSFs) of the protein’s C_α_ over the course of simulations (Figure S3) suggest that in POPC bilayers, all three forms had reached an equilibrium state and were stable. The tetrameric UCP2 showed the smallest fluctuations in RMSD and RMSF, suggesting it to be more stable than the other two forms.

**Figure 4.**
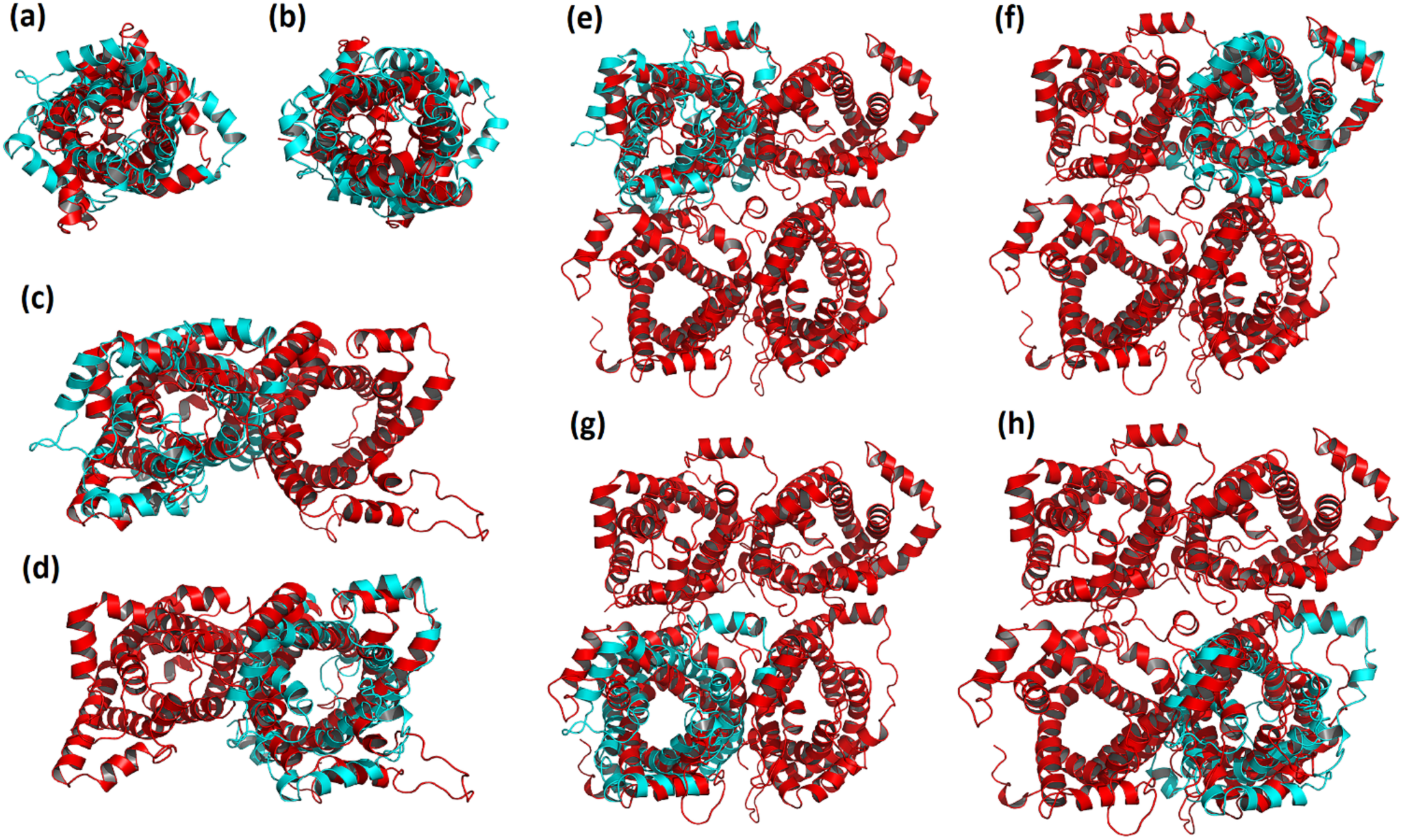
Overlay of the monomer (a-b), dimer (c-d) and tetramer (e-h) final MD structure (red) with the initial structure (PDB ID: 2LCK, cyan). An overall shrinkage is observed upon insertion of the protein into the POPC bilayer. The monomeric subunits within the dimeric and tetrameric structures are not identical, leading to asymmetric protein oligomers.

Figure 4 reveals that despite the general resemblance between the initial and final MD structures, and among different multimeric states, there are subtle yet notable, changes that occur upon insertion of the protein into the bilayer. Generally, all three forms of UCP2 underwent compaction over the course of simulation. In the case of the monomeric UCP2, the TM helix 5 (residues 214-243) was shortened by about two turns during the simulation (residues 222-243), elongating the linker between TM helices 4 and 5 (Figure S4). Figure 4 also shows that both dimeric and tetrameric UCP2 adopted asymmetric conformations (the deviations from the initial structure were not similar among their constituting subunits). More detailed comparisons of the initial and final structures of the dimeric and tetrameric forms of UCP2 are provided in Figures S5 and S6. The differences between the monomeric and oligomeric forms of the protein arise from the changes in the local environment of the monomeric UCP2 upon association, altering the modes of protein-environment interactions and protein motions in the lipid bilayer. The effect of the environment on the protein motion was further explored with principal component analysis (PCA) of the proteins (Figure S7 and S8), which indicated greater motions in the TM helices for the multimeric UCP2 in comparison to the monomeric protein during the course of simulation. As seen in Figure S8, certain parts of the subunits of dimer and tetramer moved in different manners resulting in the overall contraction of the protein. This contraction was especially observable in the tetramer. Based on the PCA results (Figure S8), TM helices tended to come closer to each other in the tetramer to form a compact structure. Acquiring more compact structures was the proteins’ response to the surrounding lipid environment, minimizing the exposure of their polar residues to the hydrophobic POPC hydrocarbon tails and/or to the non-polar residues of the neighboring protein(s). The contraction of the protein structure resulted in reduction of its radius of gyration (Rg) and solvent accessible surface area (SASA) over time (Figure S9). Furthermore, the loops on both matrix and cytoplasmic sides of the protein were able to move more freely throughout the simulation and might form transient hydrogen bonds with one another and solvent, keeping the multimeric structures intact and protecting the transport controller gates. These gates (salt-bridge networks) were proposed to regulate the transport in MCF carriers (79).

In agreement with a previous all-atom MD study of monomeric UCP2 (25), the SASA of all three molecular forms of UCP2 (Figure S9) were considerably reduced in the lipid bilayer compared to the initial structure (monomeric) in detergent micelles. Variation of protein SASA in different environments seems to be common and has been reported for a number of proteins (49-51). As seen in Figure S9b, SASA of the tetramer was the smallest among the considered UCP2 forms.

Multimerization of UCP2 slightly impacted the interaction of the protein with its neighboring phospholipids. The order parameters of these phospholipids decreased as the number of protein subunits increased in the order of monomer> dimer> tetramer) (Figure S10). As seen in Figure 5, clusters of lipids at the interfaces of neighboring protein subunits act as bridges to stabilize the oligomeric form. Different motion rates of interface and bulk lipids support this bridging role (Movie S1).

**Figure 5.**
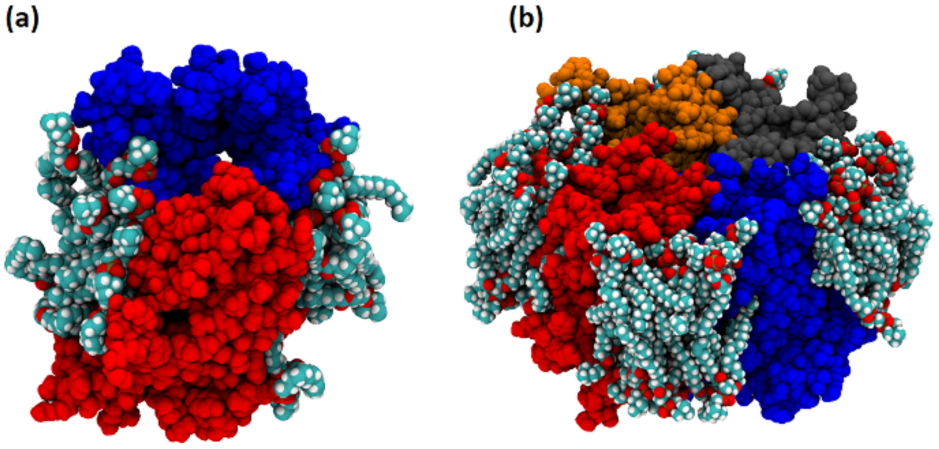
Bridging lipids in (a) dimeric and (b) tetrameric protein systems can support the stability of UCP2 oligomeric structures. The monomeric subunits of UCP2 are shown in blue, red, orange and grey. The POPC lipid molecules are colored by elements; light blue for carbon, white for hydrogen and red for oxygen atoms.

In addition to hydrophobic interactions between the protein’s helices and the bilayer, hydrogen bonds (H-bonds) also contribute to the stabilization of the protein. Analysis of different molecular forms of UCP2 showed that in the case of monomer on average 226 inter- and intra-helical and 32 protein-lipid head group H-bonds (d(donor**···**acceptor) ≤ 0.35 nm and ∠(acceptor–donor–hydrogen) ≤ 30°) were formed. These numbers increased to the respective values of 436 and 58 in the case of dimer. For tetramer the number of inter- and intra-helical and protein-lipid head group H-bonds rose to 872 and 72, respectively (Figure S11).

Salt-bridges were also responsible for stabilizing and maintaining the multimers. Interestingly, stabilizing salt bridges between the monomeric subunits (A and B) of the dimer (K109···D202, E112···K206 and K164···E46), could be observed in the same domains between the monomeric subunits of dimers A-B (K109···D202, K164···E264 and E306···K206) and C-D (R155···E46, K164···E46 and E306···K206) within the tetramer (Table 1 and Figure S12). The flexibility of the loops and the fact that there were multiple charged residues on the loops, warrant the formation of salt-bridges between the monomeric subunits. On the other hand, no salt-bridges were formed between the two dimers. This suggests that the tetrameric form is in fact a dimer of dimers and, in comparison to dimers formed of monomers, these dimers are loosely bound. Moreover, umbrella sampling simulations revealed a smaller dissociation energy for the tetramer into two dimers (60.2 kcal/mol) compared to the dissociation of the dimer (78.7 kcal/mol; Figure 6). These observations are consistent with our previous experimental reports on UCPs (3, 22) (see Discussion section).

**Table 1.** Salt-bridges formed in the multimeric UCP2 systems.^**a**,**b**^

|  |  |  |
| --- | --- | --- |
| Dimer | Subunit A···Subunit B |  |
|  | K109 | D202 |
|  | E112 | K206 |
|  | K164 | E46 |
| Tetramer | Subunit A···Subunit B |  |
|  | K109 | D202 |
|  | K164 | E264 |
|  | E306 | K206 |
|  | Subunit C···Subunit D |  |
|  | R155 | E46 |
|  | K164 | E46 |
|  | E306 | K206 |

<sup>a</sup> Subunits A, B, C and D correspond to the blue, red, orange and grey monomeric units shown in Figure 6, respectively. <sup>b</sup> Salt-bridges are ordered based on the number of subunit A (or C) amino acids involved.

**Figure 6.**
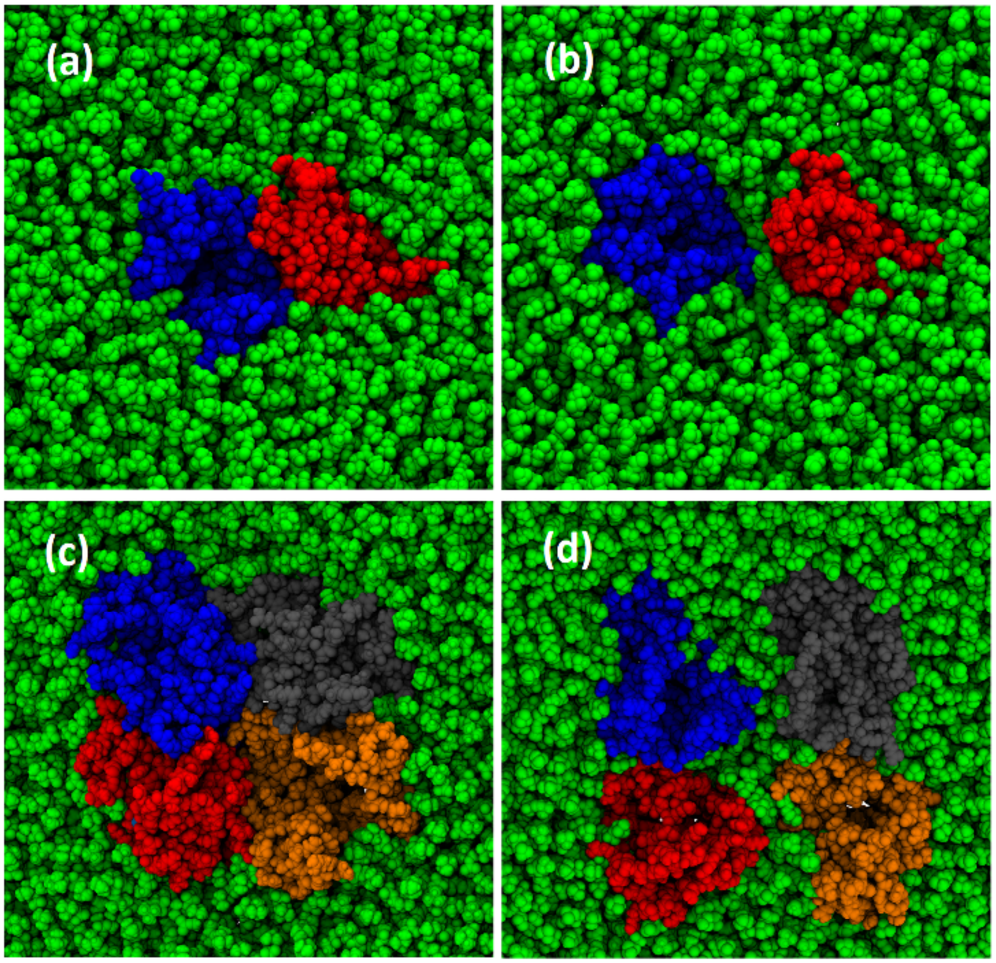
Initial and final structures of the dimeric (a-b) and tetrameric (c-d) UCP2 used for the dissociation process. Dissociation of dimer to its constituting monomers needs higher energy (18.5 kcal more) than dissociation of tetramer to its constituting dimers, suggesting tighter packing of monomers within the dimer compared to the two dimers within the tetramer. Subunits A, B, C and D are shown in blue, red, orange and grey, respectively.

As expected, oligomerization affected the lateral diffusion of the protein. The mean square displacement (MSD) of the protein reveals a decrease with oligomerization, as shown in Figure S13. The smaller lateral diffusion of tetrameric UCP2 restricts further aggregation of tetramers. Figure 7 demonstrates how different association states of UCP2 can affect the charge density (a-c) and electric potential (d-f) of the bilayer-protein system. Several computational studies have previously examined such properties across other membrane proteins (80) and lipid bilayers by exerting external electric fields (81-83) or ion imbalance on the two sides of the membrane (81, 84). A comparison of proteins’ charge density in monomeric (a), dimeric (b) and tetrameric (c) forms indicates that compared to the monomer, the magnitude of charge density did not change linearly upon oligomerization. However, despite the changes in the components’ contributions, the overall charge density and the spatial charge distribution of the systems has not remarkably changed upon oligomerization (Figure 7g).

**Figure 7.**
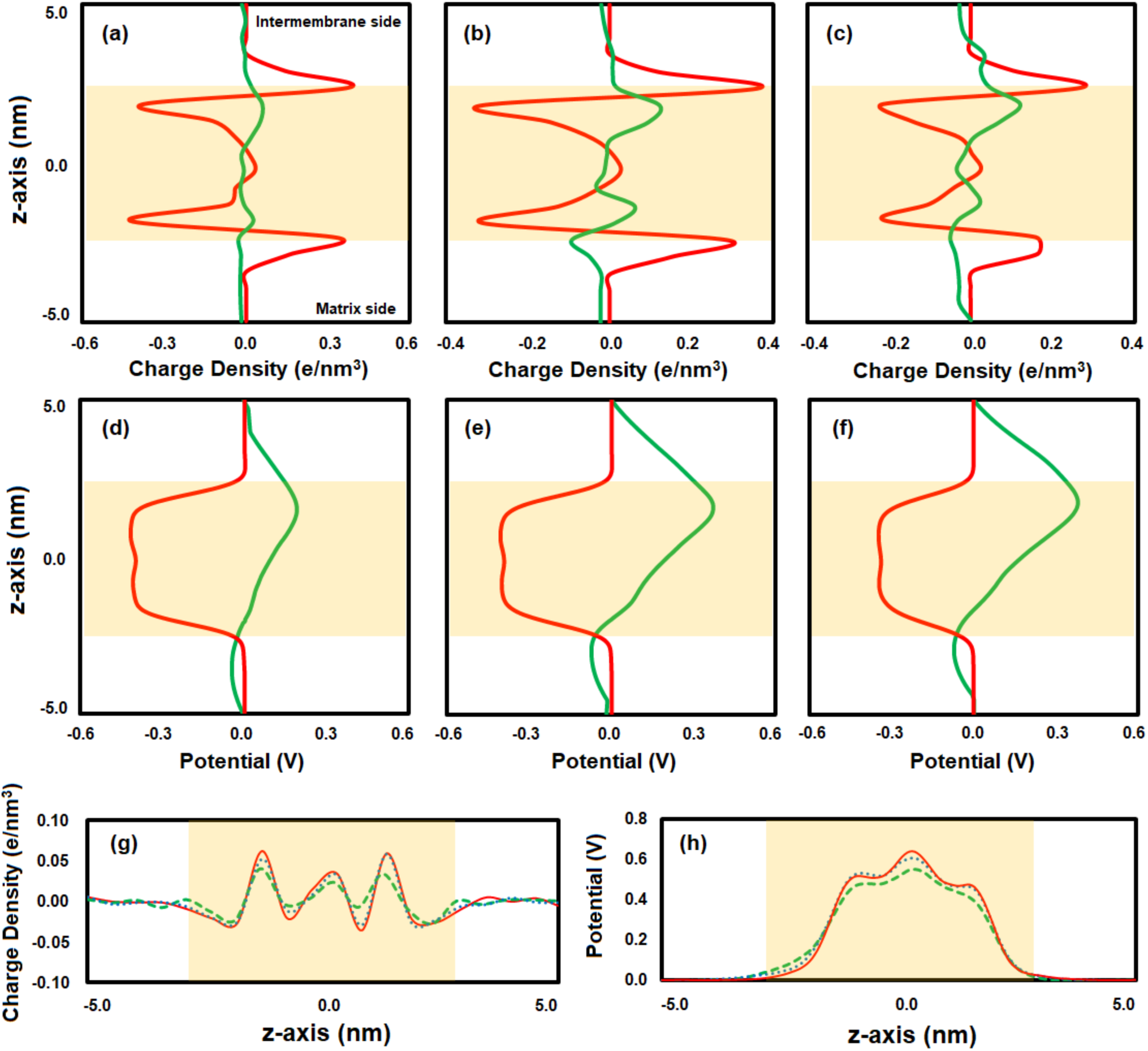
Individual contributions to the charge density (bilayer (red) and protein (green)) for (a) monomeric, (b) dimeric, and (c) tetrameric UCP2 along z-axis. Electrostatic potential contributions (bilayer (red) and protein (green)) in the (d) monomeric, (e) dimeric, and (f) tetrameric UCP2 along z-axis. (g) Total charge density and (h) total electrostatic potential of the system for the monomeric (red), dimeric (dotted blue), and tetrameric (dashed green) UCP2. The approximate position of the bilayer is shown in cream. The ionic strength was 0.15 M KCl, and no external electric field was applied.

The electrostatic potential of all three molecular states of the protein was more positive on the intermembrane space side of the bilayer (Figure 7, d-f). This suggests that the general electric properties of UCP2, together with the external electrochemical potential across the mitochondrial inner membrane, might facilitate the proton transport and selective passage of the positive ions toward the matrix side via a specific intramolecular mechanism. In comparison to the monomer, the electrostatic potential doubled upon dimerization (7d and 7e). Nevertheless, the electrostatic potential value did not change significantly upon tetramerization from dimers (7f). The nonlinear increase in the potential and charge density from monomeric to tetrameric form indicates an asymmetric arrangement of the subunits in the multimeric forms. The asymmetric electrostatic potential of UCP2 (d-f) is consistent with the role of the protein as a transporter. Furthermore, compared to monomeric and dimeric forms, the protein-bilayer system potential for the tetramer is decreased, implying a more stable molecular arrangement (Figure 7h).

## DISCUSSION

Several functions, including proton motive force attenuation and ROS regulation, have been proposed for UCP2 (6-8). On the other hand, the only published atomic structure of any UCP is that of UCP2, using a semi-empirical fragment searching NMR analysis (16). The functional molecular form of UCPs (in particular UCP1) in mitochondria has been a subject of debate, with some researchers proposing monomers (85) and others proposing oligomers as possible functional forms of the protein (3, 19, 22).

The molecular characterization of the functional structure(s) of UCPs is essential for understanding their mechanisms of physiological action.

In the current study, it has been experimentally shown that UCP2 spontaneously self-associates to form tetramers (M_w_ ∼132 kDa) in lipid membranes (Figure 1). This tetrameric UCP2 is capable of transporting protons (Figure 2), a general characteristic of all UCPs (3, 19, 22), On the other hand, AAC1, a mitochondrial carrier protein with comparable three-dimensional structure to UCP2, does not form tetramers in lipid membranes. These observations indicate the specific tendency of UCP2 (and other UCPs, as shown previously) to form tetramers in phospholipid bilayers, which cannot be generalized to all MCF members. UCP2’s tetrameric complex is in fact a dimer of dimers, in which the interactions between the monomers forming the dimer are stronger than the interactions between the dimers within the tetramer. The specific tendency of UCP2 to associate in lipids and the interaction between monomeric and dimeric units in the tetrameric protein, are further supported by MD simulations. Physicochemical evidence for the formation of associated UCPs, including analytical ultracentrifugation (sedimentations equilibrium measurements) and mass spectrometric analyses, have been previously reported (3, 22). Overall, the experimental and computational results of the current study on association of UCP2 and stability of its tetrameric state are consistent with our previous studies on other UCPs (3, 22).

In addition to the formation and stability of the tetrameric protein, confirmed both in the current and previous (3) studies, the reversal of the ellipticity ratio (θ_208_/θ_222_) after reconstitution of the monomeric proteins (from OG detergent) into liposomes (Figure 3a), implies both a more compact interhelical packing of monomers as well as stronger intermolecular interactions between these monomers (3, 22). The low rate of proton transport and inhibition of this transport by common UCP inhibitors such as ATP (for both reconstituted monomer and monomer/multimer mixture), also confirms that the tetrameric UCP2 is functional in lipid membranes, and is more likely to be a transporter than a channel.

The MD simulations of the tetramer in bilayers show that despite the compaction of all constituting monomers over the course of simulation (comparable to changes in the ellipticity ratio mentioned in the previous paragraph), these subunits do not adopt similar structures and lateral orientations (Figure 4). This asymmetric conformation of monomeric units in the tetramer can also be observed in the electric behavior of the protein and may play a role in the electrochemical regulation of its ion transport function. The asymmetric nature of the tetramer could provide a scaffold for regulating the activity of individual units, and also facilitate the cooperative intercommunication between these subunits. Concomitantly, all individual monomeric subunits might not be active at the same time. Asymmetric oligomerization has also been reported for other proteins such as tetrameric D-Glyceraldehyde-3-phosphate dehydrogenase (86) and dimeric Tryptophanyl-tRNA synthetase (87).

Detailed analysis of the interactions stabilizing the dimeric and tetrameric UCP2 indicates that comparable salt-bridges are formed between the monomers within the dimer and between the monomers within the dimeric units constituting the tetramer (Table 1). No salt-bridges are detectable between the two dimeric units of the tetramer, implying that, in comparison to monomers, the dimers are less strongly bound. The specific electrostatic interaction between the monomeric, rather than dimeric, units supports our original conclusion that tetrameric UCPs are indeed dimer of dimers (Figure 6) (22). This hypothesis is further supported by the higher dissociation energy of monomeric subunits of a dimer compared to dimeric subunits of a tetramer by 18.5 kcal. The computational results of this study are also in agreement with our previous report, in which titration of tetrameric UCP2 using an anionic detergent (SDS) resulted in dissociation to dimers at low SDS concentrations (slightly above detergent’s CMC, ≳ 2 mM) and monomers at high SDS concentrations (≳ 83 mM) (3).

In addition to electrostatic and other non-covalent protein-protein interactions, the tetrameric assembly of UCP2 in lipid bilayers can be also stabilized by bridging lipids (Figure 5 and movie S1). Since tetramerization of the UCP family in model lipid bilayers has been observed in PC/PE, POPC and POPC/cardiolipin vesicles (3, 22), it can be suggested that the role of lipids in UCP’s self-association is non-specific. However, further experiments are required for verifying the role of specific lipids in the stability and ion transport function of UCPs; the role of lipids cannot be easily generalized as has been shown in the case ATP binding cassette proteins where results have shown both dependence and independence of lipid specificity depending on the systems and properties of interest (88-90). The importance of non-specific protein-lipid interaction is also exhibited in the increased helicity of the protein in lipid bilayers (Figure 3a). The role of bridging lipids in stabilizing the multimeric complexes of membrane proteins has been shown for several proteins such as dimeric *Aquifex aeolicus* leucine transporter, dimeric *E. coli* Na^+^/H^+^ antiporter and tetrameric aquaporin (aqpZ) (34).

## CONCLUSION

In conclusion, detailed structural and ion transport studies of UCP2 in lipid bilayers, as well as computational simulations of the dynamics of UCP2 oligomerization in membranes, were utilized to provide a deeper molecular insight into the structure and function of uncoupling proteins. The findings of this study also implicate the possibility of simultaneous co-existence of functional monomeric and dimeric, in addition to tetrameric, protein under various cellular conditions and different membrane lipid compositions. However, the scope of *in vitro* studies is always limited by the use of detergents in protein purification and reconstitution steps, and by not being able to mimic the exact dynamic physicochemical properties of the IMM lipid/protein membranes. These limitations can lead to deviations from the structure and function of proteins in living cells, and the real-time *in-vivo* analysis of IMM, considering the complexities involved, is still to be anticipated.

## SUPPLEMENTAL INFORMATION

The following Supplemental Information can be found online at https://doi.org/…

-Representation of oligomerization motifs on UCP2 structure

-Western blot analysis of AAC1

-Backbone RMSD and Cα RMSF of UCP2

-Overlay of monomer and 2LCK

-Overlay of dimer and 2LCK

-Overlay of tetramer and 2LCK

-Projection of the MD simulation trajectories on the first 4 eigenvectors

-Visualization of principle component analysis

-Radius of gyration and solvent accessible surface are of UCP2

-Lipid order parameter

-Number of hydrogen bonds over time

-Salt-bridges between the neighboring UCP2 units

-Mean square displacement of UCP2

-Movie clip of specific interaction of lipids with the UCP2 tetrameric unit

## AUTHOR CONTRIBUTIONS

A. A. performed all protein expression, reconstitution, spectroscopic and proton transport experiments for UCP2, as well as drafting the article. S.O. U. initiated and performed the preliminary work on UCP2 expression and purification. M. F. assisted with and performed protein cloning, expression, reconstitution and spectroscopic experiments for AAC1. S. S.-H. planned and performed the molecular dynamics simulation and computational work for UCP2, and drafted the related sections. M. K. planned and provided technical advice on the computational section, as well as the entire text. M.D. S. provided technical advice and expertise with the molecular cloning and expression of proteins, and the entire text. M. J.-N. conceived the study and participated in data analysis and drafting of the article.

## ACKNOWLEDGMENT

This research project was supported by Natural Sciences and Engineering Research Council of Canada (NSERC) discovery and Canada Foundation for Innovation (CFI) grants to MJ-N (05900 and 6786), MDS (05437 and 11292), and MK. MK thanks the Canada Research Chairs Program. AA is a recipient of Ontario Trillium Scholarship. Computing facilities have been provided by SHARCNET (www.sharcnet.ca) and Compute Canada (www.computecanada.ca).

