## Supporting Information for "Experimental and Computational Evidence for Self-Assembly of Mitochondrial UCP2 in Lipid Bilayers"

**Afshan Ardalan[1], Stephanie O. Uwumarenogie[1], Michael Fish[1,2], Shahin Sowlati-Hashjin[3,4], Mikko Karttunen[3,4,5], Matthew D. Smith[2], and Masoud Jelokhani-Niaraki[1]***

[1] Department of Chemistry and Biochemistry, Wilfrid Laurier University, Waterloo, ON, Canada N2L 3C5; [2] Department of Biology, Wilfrid Laurier University, Waterloo, ON, Canada N2L 3C5; [3] Department of Chemistry, University of Western Ontario, London, ON, Canada N6A 3K7; [4] The Center for Advanced Materials and Biomaterials Research, The University of Western Ontario, London, Ontario, Canada N6K 3K7; [5] Department of Applied Mathematics, Western University, London, ON, Canada N6A 5B7

**Table of Contents**

| | | |
|---|---|---|
| **Figure S1.** | Representation of oligomerization motifs on UCP2 structure | S2 |
| **Figure S2.** | Western blot analysis of AAC1 | S2 |
| **Figure S3.** | Backbone RMSD and Cα RMSF of UCP2 | S3 |
| **Figure S4.** | Overlay of monomer and 2LCK | S4 |
| **Figure S5.** | Overlay of dimer and 2LCK | S5 |
| **Figure S6.** | Overlay of tetramer and 2LCK | S6 |
| **Figure S7.** | Projection of the MD simulation trajectories on the first 4 eigenvectors | S6 |
| **Figure S8.** | Visualization of principle component analysis | S7 |
| **Figure S9.** | Radius of gyration and solvent accessible surface are of UCP2 | S8 |
| **Figure S10.** | Lipid order parameter | S9 |
| **Figure S11.** | Number of hydrogen bonds over time | S10 |
| **Figure S12.** | Salt-bridges between the neighboring UCP2 units | S11 |
| **Figure S13.** | Mean square displacement of UCP2 | S11 |

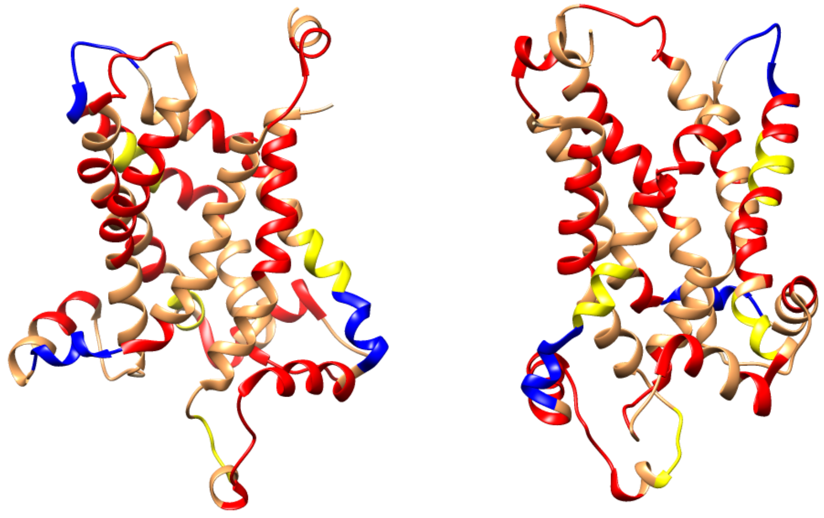

**Figure S1.** The two interacting faces of UCP2 selected to form a dimer. The GXXXG, GXXXLXXG, and SmXXXSm (Sm = Gly, Ala, Ser, Thr) motifs, which promote oligomerization are shown in yellow, blue, and red, respectively.

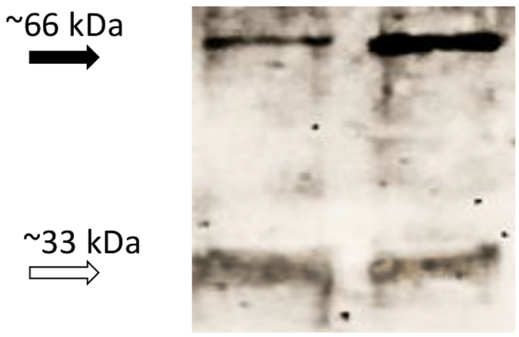


**Figure S2.** Western blot analysis of AAC1 probed with α-AAC1 antibody confirms the co-presence of monomeric and dimeric states in OG.

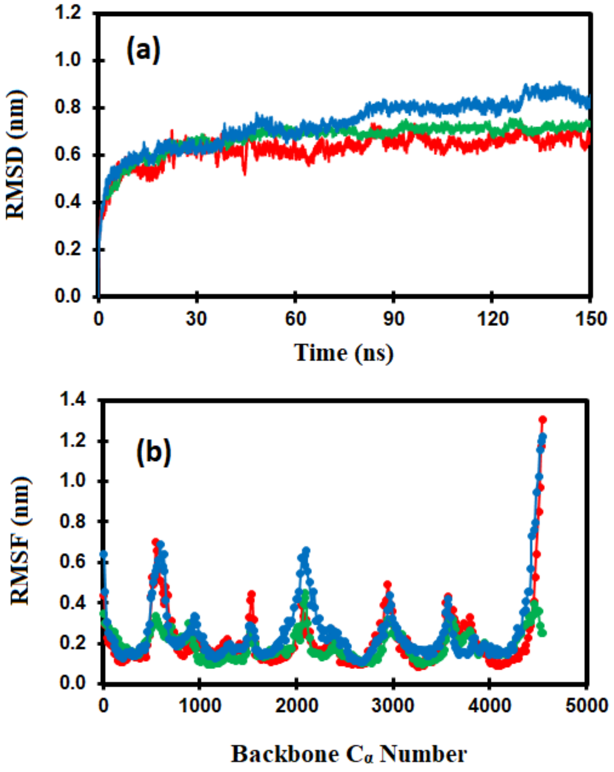


**Figure S3.** (a) Backbone RMSD of monomeric (red), dimeric (blue), and tetrameric (green) UCP2 in the membrane over the course of MD simulation. (b) RMSF of $C_\alpha$ of protein for monomeric (red), dimeric (blue), and tetrameric (green) UCP2 in the membrane over the course of MD simulation.

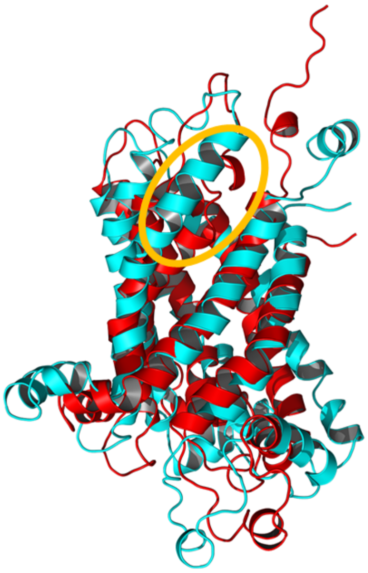

**Figure S4.** Overlay of the monomer final MD structure (red) with the initial structure (PDB ID: 2LCK, cyan) along x-axis. The shortened helix 5 during simulation is circled in yellow. This helix is shortened over the course of simulation leading to an increase in the length of the loop between helices 4 and 5.

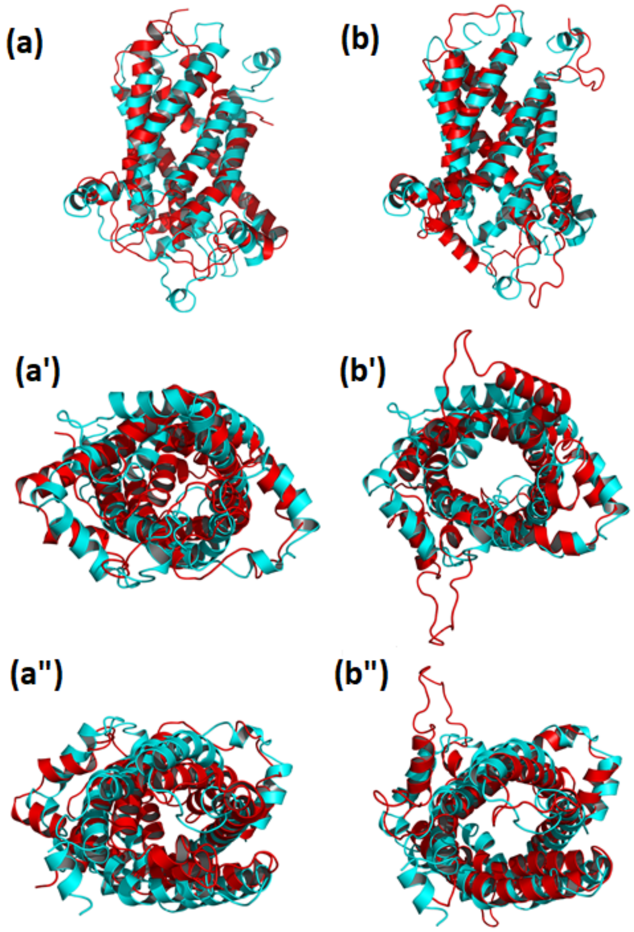


**Figure S5.** Overlay of the final MD structure of dimer units (red) with the initial structure (PDB ID: 2LCK, cyan) along (a – b) x-axis, (a′ – b′) z-axis (top view), and (c – c′) z-axis (bottom view).

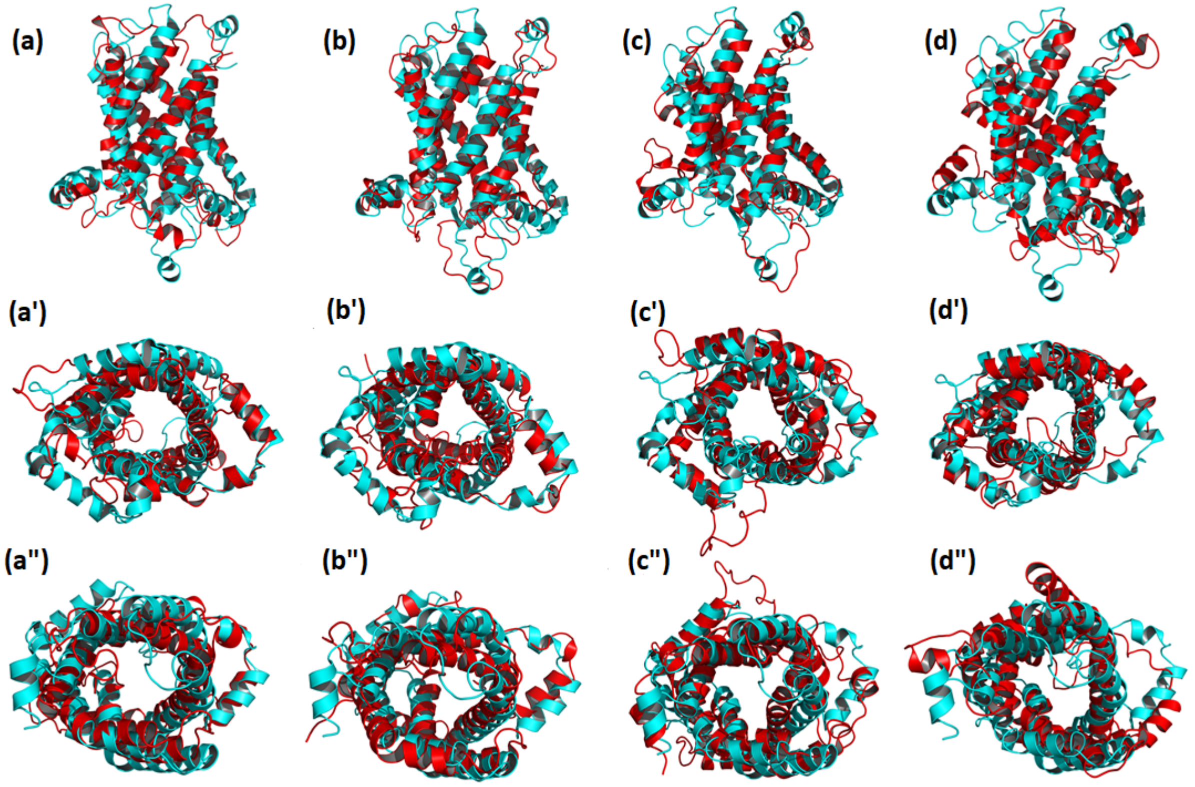


**Figure S6.** Overlay of the final MD structure of tetramer units (red) with the initial structure (PDB ID: 2LCK, cyan) along (a - d) x-axis, (a′ - d′) z-axis (top view), and (a″ - d″) z-axis (bottom view).

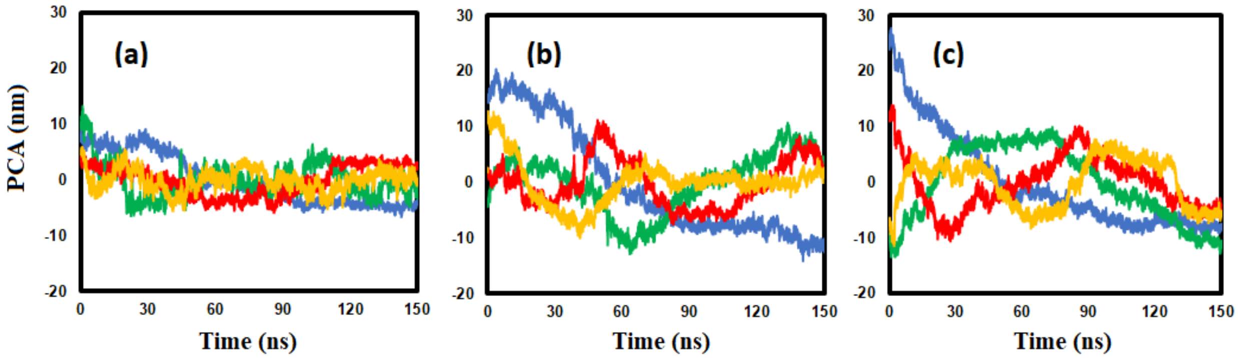


**Figure S7.** Projection of the MD simulation trajectories on the first 4 (1st (blue), 2nd (green), 3rd (red), 4th (yellow)) eigenvectors (principal component, PC) for (a) monomeric, (b) dimeric, and (c) tetrameric UCP2 in POPC bilayer.

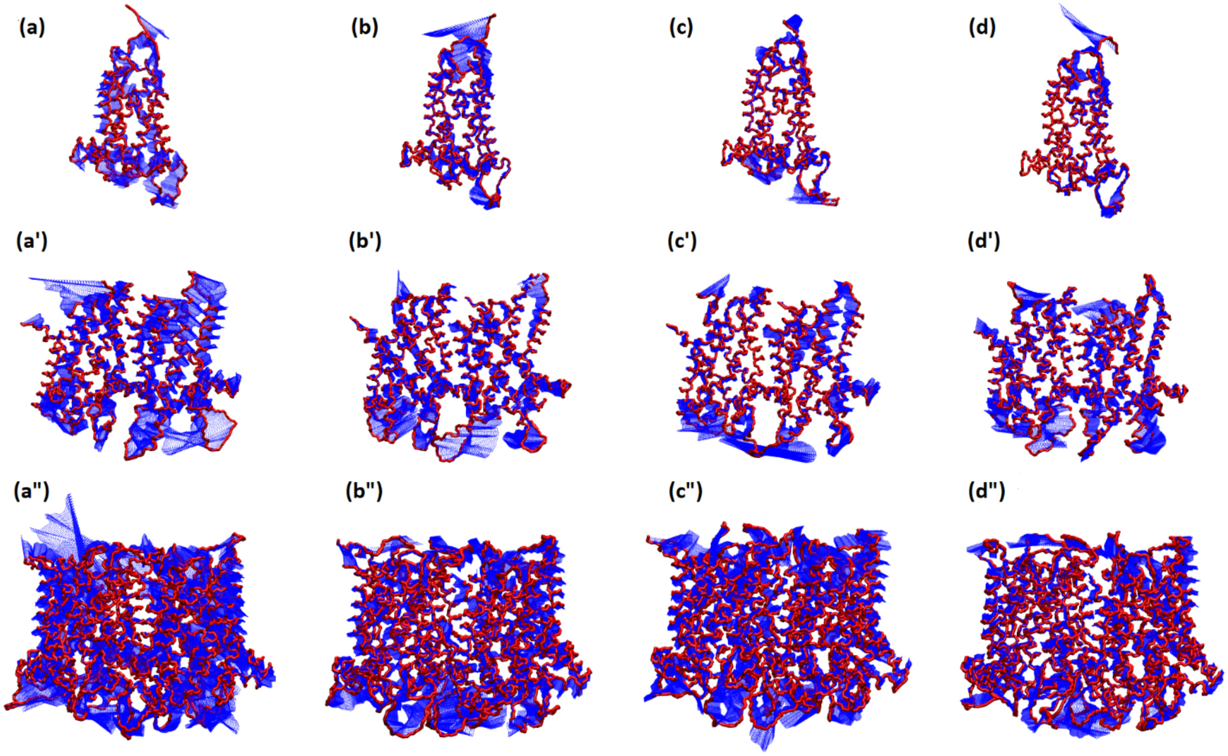


**Figure S8.** Visualization of PCA: Projections of the MD simulations on the first four eigenvectors for the monomeric (a – d), dimeric (a′ – d′), and tetrameric (a″ – d″). Blue hatching represents the direction of motions.

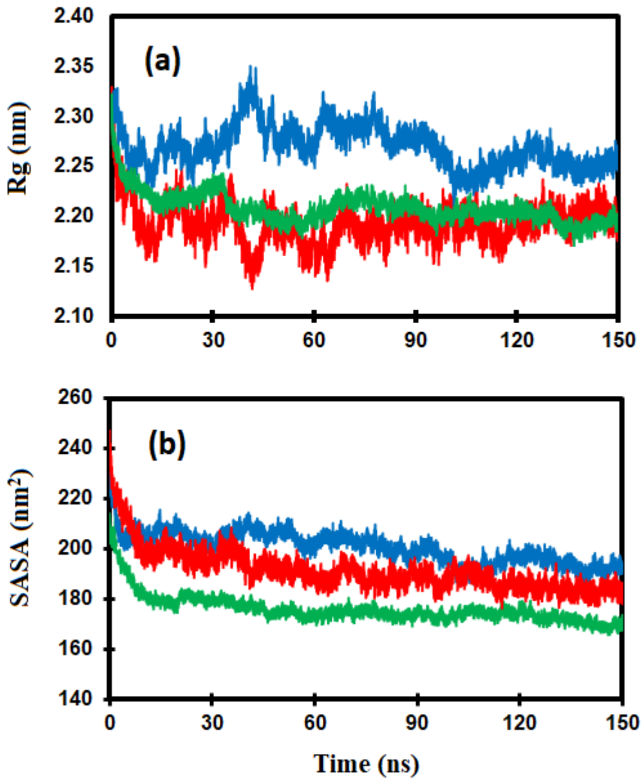


**Figure S9.** (a) Radius of gyration for the monomeric (red), dimeric (blue), and tetrameric (green) UCP2 in the membrane over the course of MD simulation. (b) Solvent accessible surface area (SASA) of monomeric (red), dimeric (blue), and tetrameric (green) UCP2 in the membrane over the course of MD simulation.

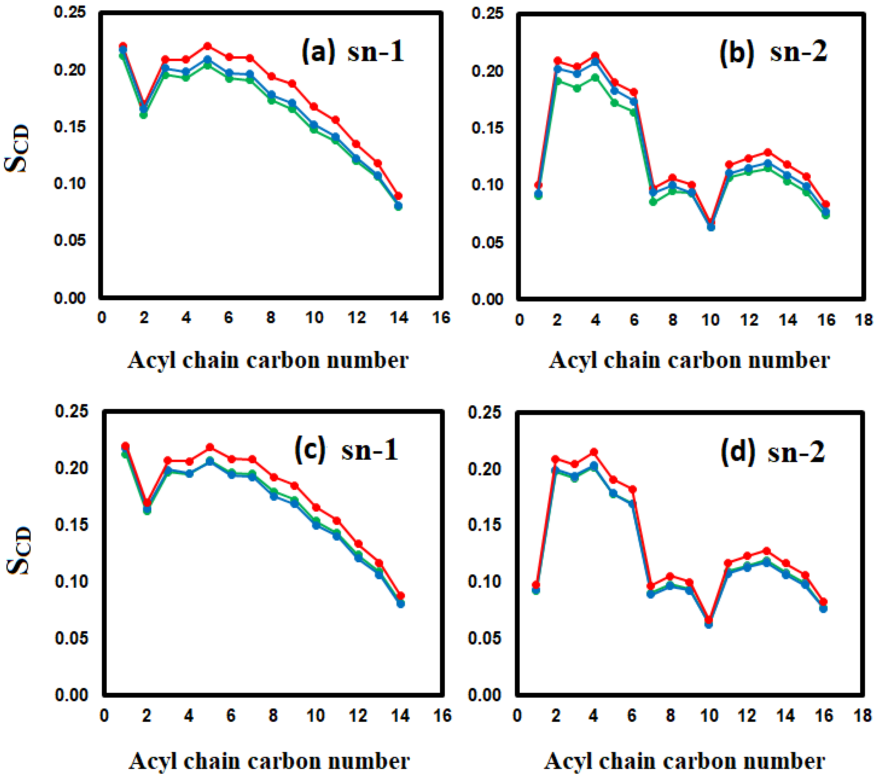


**Figure S10.** Order parameter for (a) chain 1 (sn-1) and (b) chain 2 (sn-2) of all phospholipids, (c) chain 1 (sn-1) of only protein-neighboring and (d) chain 2 (sn-2) of only protein-neighboring phospholipids for monomeric (red), dimeric (blue), and tetrameric (green) UCP2. The order parameter, $S_{CD}$ is defined as $\langle(3\ cos\theta_n - 1)/2\rangle$, where $\theta_n$ is the angle between nth segmental vector along the acyl chain and the phospholipid bilayer normal.

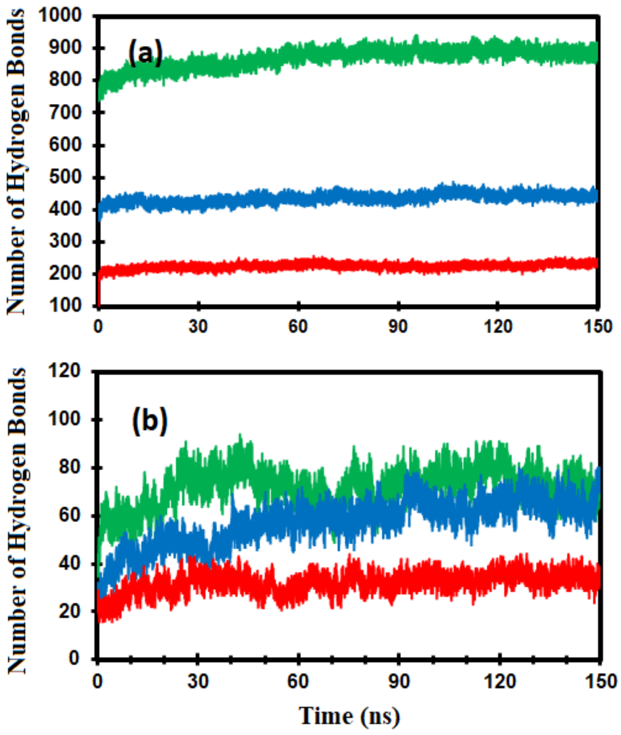


**Figure S11.** (a) Number of inter- and intra-helical hydrogen bonds with time for (red) monomeric, (blue) dimeric, and (green) tetrameric UCP2. (b) Number of hydrogen bonds between (red) monomeric, (blue) dimeric, and (green) tetrameric UCP2 and POPC bilayer.

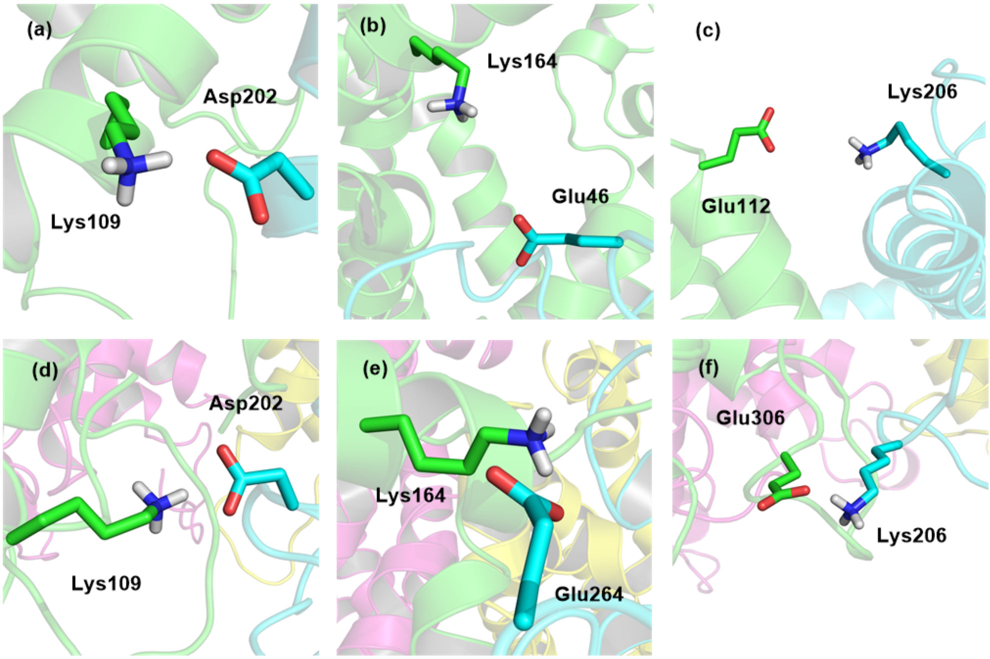


**Figure S12.** Main salt-bridges between the neighboring UCP2 units in dimeric (a – c) and in tetrameric (d – f) forms. Subunits A, B, C, and D are shown in green, cyan, purple, and yellow, respectively.

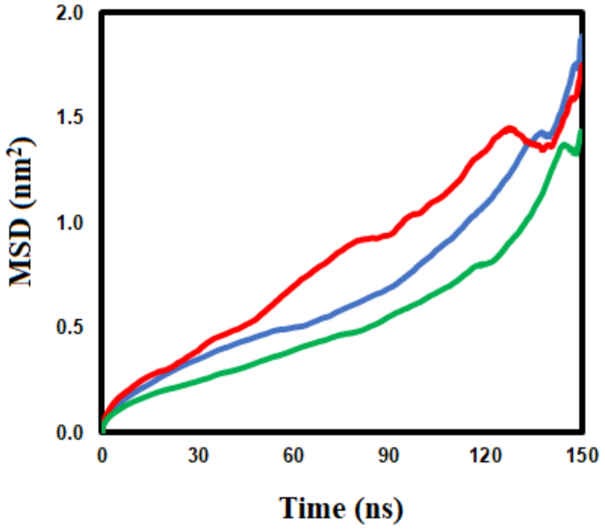


**Figure S13.** Mean square displacement of monomeric (red), dimer (blue), and tetrameric (green) UCP2 in POPC bilayer with time.
